# Insulinoma-associated 1 promotes neurogenic proliferation of cortical basal progenitors but is largely dispensable for projection neuron production

**DOI:** 10.64898/2026.04.02.716139

**Authors:** Venkata Thulabandu, Xinwei Cao

## Abstract

Basal progenitors (BPs) are essential contributors to mammalian cortical neurogenesis, yet the mechanisms governing their behavior remain incompletely understood. Insulinoma-associated 1 (INSM1), a SNAG-domain zinc-finger transcription factor, has been proposed to promote BP generation and expansion, although prior conclusions relied heavily on gain-of-function approaches and were limited by early lethality of *Insm1*-null embryos. Here, using conditional genetic ablation of *Insm1* in mouse cortical progenitors, we reveal that INSM1 is largely dispensable for BP generation but is essential for proper BP cell-cycle progression. INSM1-deficient BPs exhibit impaired S-phase entry, reduced RB phosphorylation, and downregulation of cell-cycle–related gene programs. Although neurogenesis by BPs is diminished, apical progenitors (APs) compensate by increasing symmetric amplifying divisions, expanding the AP pool and preserving production of later-born, upper-layer neurons despite reduced early-born, deep-layer neurons. These findings identify INSM1 as a critical regulator of BP neurogenic proliferation and highlight compensatory flexibility within the cortical progenitor hierarchy.

**Significance Statement:** The developing mammalian cortex contains two principal classes of neural progenitor cells: apical progenitors (APs), which serve as the primary stem/progenitor population; and basal progenitors (BPs), which are produced by APs. The vast majority of cortical neurons are generated by BPs, yet how BP proliferation is regulated remains unclear. We show that Insulinoma-associated 1 (INSM1), previously thought to control BP biogenesis, instead governs BP cell-cycle progression. Loss of INSM1 impairs BP proliferation. However, overall cortical neuron output is largely preserved because APs compensate by expanding their population. Our findings revise the role of INSM1 in cortical development and provide insight into how mammalian neurogenesis maintains robustness through compensatory responses among neural progenitor populations.

## Introduction

The developing vertebrate central nervous system (CNS) contains two principal classes of neural progenitor cells: apical progenitors (APs) and basal progenitors (BPs) (Taverna et al., 2014). APs, which include neuroepithelial cells and the cells they transform into after the onset of neurogenesis—apical radial glia (aRG), are the primary stem/progenitor cells. With their cell bodies forming the ventricular zone (VZ) lining the neural tube lumen, APs undergo mitosis at or very close to the ventricular surface (VS). BPs, which are produced by APs and reside in the subventricular zone (SVZ) adjacent to the VZ, undergo mitosis away from the VS.

APs are capable of multiple rounds of cell division, which can be symmetric amplifying divisions that generate two APs, or asymmetric neurogenic divisions that produce one AP together with either a neuron or a BP, and finally a terminal self-consuming division that yields two non-AP progeny. The prevalence of each division mode varies by species, anatomical region, and developmental stage. For example, in the developing mouse neocortex, APs predominantly undergo symmetric amplifying divisions during early neurogenesis and switch almost entirely to asymmetric neurogenic divisions by embryonic day (E) 13.5 (Gao et al., 2014).

While APs ultimately generate all neurons and glia, the contribution of BPs to neurogenesis differs across species and anatomical regions (Thor, 2024). BPs are thought to be particularly important for the development and evolution of the mammalian cerebral cortex (Martínez-Cerdeño et al., 2006; Molnár et al., 2006; Cheung et al., 2007; Cheung et al., 2010; Borrell and Calegari, 2014). Organized into six layers, glutamatergic projection neurons in the mammalian neocortex are produced in a general “inside-out” order in which later-born neurons migrate past earlier-born neurons to progressively occupy more superficial layers (Angevine and Sidman, 1961; Rakic, 1974). In the mouse neocortex, approximately 70% of deep-layer (layers 5–6) projection neurons and 90% of upper-layer (layers 2–4) projection neurons are generated by BPs (Huilgol et al., 2023). In gyrencephalic species, the expansion of cortical projection neurons is accompanied by an increased number and diversity of BPs, along with enhanced proliferative capacity and altered division modes (Hansen et al., 2010; Lui et al., 2011; Dehay and Huttner, 2024). Whereas the vast majority of BPs in the developing mouse neocortex undergo a single round of terminal symmetric division to produce two neurons (Haubensak et al., 2004; Miyata et al., 2004; Noctor et al., 2004; Noctor et al., 2008; Mihalas and Hevner, 2018; Huilgol et al., 2025), BPs in the developing primate neocortex—including basal/outer radial glia—can undergo multiple rounds of symmetric amplifying or asymmetric neurogenic divisions (Taverna et al., 2014; Dehay and Huttner, 2024).

Given the importance of BPs during cortical development and evolution, elucidating their molecular regulators is critical. Insulinoma-associated 1 (INSM1), a SNAG-domain zinc-finger transcription factor (Breslin et al., 2002; Lan and Breslin, 2009; Chiang and Ayyanathan, 2013), was previously proposed to function as “a master regulator of BP biogenesis” in the neocortex (Farkas et al., 2008). Forced expression of INSM1 converts APs into BPs, whereas *Insm1* deletion reportedly reduces the numbers of BPs and neurons of all cortical layers (Farkas et al., 2008; Tavano et al., 2018). These conclusions, however, have several caveats. First, they rely heavily on gain-of-function analyses. Second, because *Insm1-*null embryos die at or before E16.5, phenotypic characterization was carried out at or before E16.5, when cortical neurogenesis is still ongoing. Finally, the overall embryo health could affect neural development in a systemic, non-cell-autonomous manner. Thus, the precise role of INSM1 in cortical neurogenesis remains unresolved.

Here we conditionally deleted *Insm1* during mouse cortical development and performed comprehensive analyses of neural progenitors and projection neurons across embryonic and postnatal stages. Contrary to previous reports, we find that INSM1 is not required for BP biogenesis. Instead, INSM1 promotes BP cell-cycle progression. Furthermore, we show that although deep-layer neuron production is reduced, upper-layer neuron production proceeds normally in the absence of INSM1.

## Materials and Methods

### Mice

The *Emx1-Cre* line was obtained from the Jackson Laboratory (stock #005628) and maintained in the C57BL/6 background. *Insm1^flox/flox^*mice were kindly provided by Jaime García-Añoveros (Northwestern Feinberg School of Medicine) and maintained in a mixed background. All animal procedures were approved by the Institutional Animal Care and Use Committee of St. Jude Children’s Research Hospital (SJCRH). To label S-phase cells, EdU and BrdU were injected into timed pregnant mice intraperitoneally at 10 μg and 50 μg per gram body weight, respectively. Postnatal animals of both sexes were included as experimental subjects. The sex of embryos was not determined.

### Histology and staining

Embryonic mouse brains were dissected in cold PBS and fixed overnight in 4% paraformaldehyde (PFA) at 4°C. Postnatal animals were transcardially perfused with PBS and PFA, and their brains were dissected and fixed in PFA overnight at 4°C. For frozen sections, fixed brains were washed in PBS, equilibrated sequentially in 10% and 25% sucrose overnight at 4°C, embedded in Tissue Freezing Medium (Thermo Scientific), and sectioned at 12–20-μm thickness. For Nissl and Luxol Fast Blue staining, P21 brains were fixed in 4% PFA for 2 overnights, dehydrated in 70% ethanol overnight, then processed and embedded in paraffin. Nissl-Luxol and Hematoxylin and eosin staining was performed based on standard protocols.

### Immunofluorescence staining

Frozen sections were washed in PBS, blocked and permeabilized in PBS with 0.1% Triton X-100 (PBST) and 10% normal donkey serum for 1 h, and incubated with primary antibodies at 4°C overnight. Sections were then washed in PBST and incubated with fluorescence–conjugated secondary antibodies (Jackson ImmunoResearch Laboratories or Thermo Scientific Invitrogen) at 1:1000 dilution and DAPI for 2–3 h at room temperature. The following primary antibodies were used: Alexa Fluor 647-SOX2 (BD Biosciences, #562139, 1:100), BrdU (Abcam ab152095, 1:500), TBR2/EOMES (eBioscience, #14-4875, 1:100), phospho-Histone H3 (Ser10) (Millipore Sigma 06-570, 1:500), INSM1 (Santa Cruz Biotechnologies sc-271408 AF488, 1:100), Ki67 (Abcam ab15580, 1:100), CUX1 (Santa Cruz sc13024, 1:100), CTIP2 (Abcam ab18465, 1:500), TBR1 (Abcam ab31940, 1:100), SATB2 (Abcam ab51502, 1:100), FOXP2 (Synaptic Systems 485005, 1:200), Phospho-Rb (Ser807/811) (Cell Signaling Technologies #8516, 1:100). EdU staining was performed using the Click-iT Plus EdU kit (Thermo Fisher Scientific, C10638) according to the manufacturer’s protocol. For BrdU staining, sections were heated in 10 mM sodium citrate (pH 6.0) to 120–125°C for 2 min in a water-filled pressure cooker (Instant Pot 13-min prewarm followed by 25-min incubation) (Kearns et al., 2023).

### Image quantification

All images were acquired using a Zeiss Apotome microscope with a 20x or 40x objective lens. Regions of interest were outlined and extracted in Zeiss Zen 3.11. Nucleus segmentation was performed using the machine-learning–based segmentation method StarDist (https://github.com/stardist/stardist), which was first trained using manually annotated datasets. Fluorescent label intensity of each channel over nuclear ROIs was calculated using a custom multi-channel intensity quantification algorithm created in OMERO (https://www.openmicroscopy.org/omero/scientists/). Thresholds for signal positivity were manually determined by inspecting each image. Neocortical length was measured with Zeiss Zen 3.11 software. Data points in all quantification graphs correspond to individual animals, with each point representing the average of 2–4 sections per animal.

### Single cell RNA-seq tissue collection and library preparation

The whole later cortex of E14.5 and anterior-to-medial lateral cortex of E16.5 mouse brains were dissected and dissociated using a Neural dissociation kit (Miltenyl Biotec 130-094-802) according to the manufacturer’s protocol. Tissues were incubated in a papain-based solution for 10 min at 37°C with occasional mixing, filtered through a 40-μm strainer (pluriSelect 43-10040-40), and spun down at 800 x g for 3 min. Cells were fixed using the Parse Biosciences Evercode™ Cell Fixation kit (ECFC3300) according to the manufacturer’s protocol, slow-frozen in Mr.Frosty, and stored at −80°C for up to 2 months. Sequencing libraries were generated using the Parse Biosciences Evercode™ WT kit (ECWT3300) according to the manufacturer’s protocol, combining 3 *Insm1* cKO and 3 control samples from each stage. Libraries were sequenced on an Illumina Novaseq 6000.

### Single-cell RNA-seq data analysis

FASTQ files were uploaded to the Parse Biosciences Trailmaker server (https://www.parsebiosciences.com/data-analysis/) for quality control, filtering, clustering, and dimensionality reduction. In brief, cells with mitochondrial content <5%, fewer than 1200 transcripts detected, or flagged as doublets were filtered out. The data were then log-normalized and scaled. Dimensionality reduction was performed without data integration, using 2,500 highly variable genes (HVGs) and 30 PCs (capturing 93.19% of the data). The obtained Seurat object was further analyzed in RStudio. The Seurat object was split into E14.5_object and E16.5_object. A resolution of 0.1 was used to cluster E14.5_object, including both control and cKO cells, after dimensionality reduction (25 PCs) based on 2,500 HVGs, without regressing out cell-cycle effects. Similarly, E14.5_control_object was extracted and clustered at a resolution of 0.3 after dimensionality reduction (20 PCs) based on 2,500 highly variable features without cell-cycle regression. The E16.5_object UAMP was generated without cell-cycle regression, but the cluster labels were identified at 0.35 resolution with cell-cycle regression, as the separation between apical and basal progenitor clusters was obscured by cell-cycle–related variation. Cell-cycle phase was inferred using the *CellCycleScoring* command. Gene signature scores were calculated using the *AddModuleScore* function based on published gene lists (Bedogni and Hevner, 2021).

For Gene Set Enrichment Analysis (GSEA) using the pseudo-bulk approach, differential expression between cKO and control cells was calculated as pseudo-bulk within each annotated cell type using wilcoxauc from the presto_v1.0.0 package (Korsunsky et al., 2019) for all the protein-coding genes. Genes were ranked based on their log_2_fold-change values and altered pathways were identified by pre-ranked GSEA (Mootha et al., 2003; Subramanian et al., 2005) using the Hallmark and GOBP gene set collections with 1,000 permutations and No_collapse option. Single Cell Pathway Analysis (SCPA) was performed using SCPA’s *compare_seurat* function (Bibby et al., 2022).

### Experimental design and statistical analysis

Graphs were generated using GraphPad Prism version 11. Individual animals were considered as biological replicates (*n*). Statistical significance was assessed by unpaired two-tailed *t*-test. The threshold for statistical significance was defined as *P* < 0.05. All values are mean ± SEM (standard error of the mean). The number of animals used (*n*), statistical test used, and *P* values are indicated in figures or figure legends.

### Data and code availability

The raw single-cell RNA sequencing datasets generated in this study have been deposited in the GEO repository databased under the accession code GSExxxxxx. The code for the scRNA-seq analysis presented in this manuscript is available publicly on GitHub.

## Results

### INSM1 loss during cortical development reduces deep-layer neuron production but spares upper-layer neurons

To investigate the role of INSM1 in cortical development, we conditionally deleted *Insm1* by crossing *Insm1^flox/flox^* mice (Wiwatpanit et al., 2018) with the *Emx1^IRES-Cre^* (*Emx1-Cre*) line, which drives Cre expression selectively in cortical progenitors starting around E9.5 (Gorski et al., 2002). We performed immunostaining to confirm the depletion of INSM1 protein. At E12.5, INSM1 was expressed sporadically in cells within the VZ, most of which coexpressed the BP marker TBR2/EOMES (Englund et al., 2005) (Fig. 1A), consistent with previous reports (Duggan et al., 2008; Farkas et al., 2008; Tavano et al., 2018). In the cortex of *Insm1* conditional knockout (cKO, *Insm1^flox/flox^;Emx1-Cre*) embryos, INSM1 signal was absent, confirming efficient loss of INSM1 protein (Fig. 1B).

**Figure 1.**
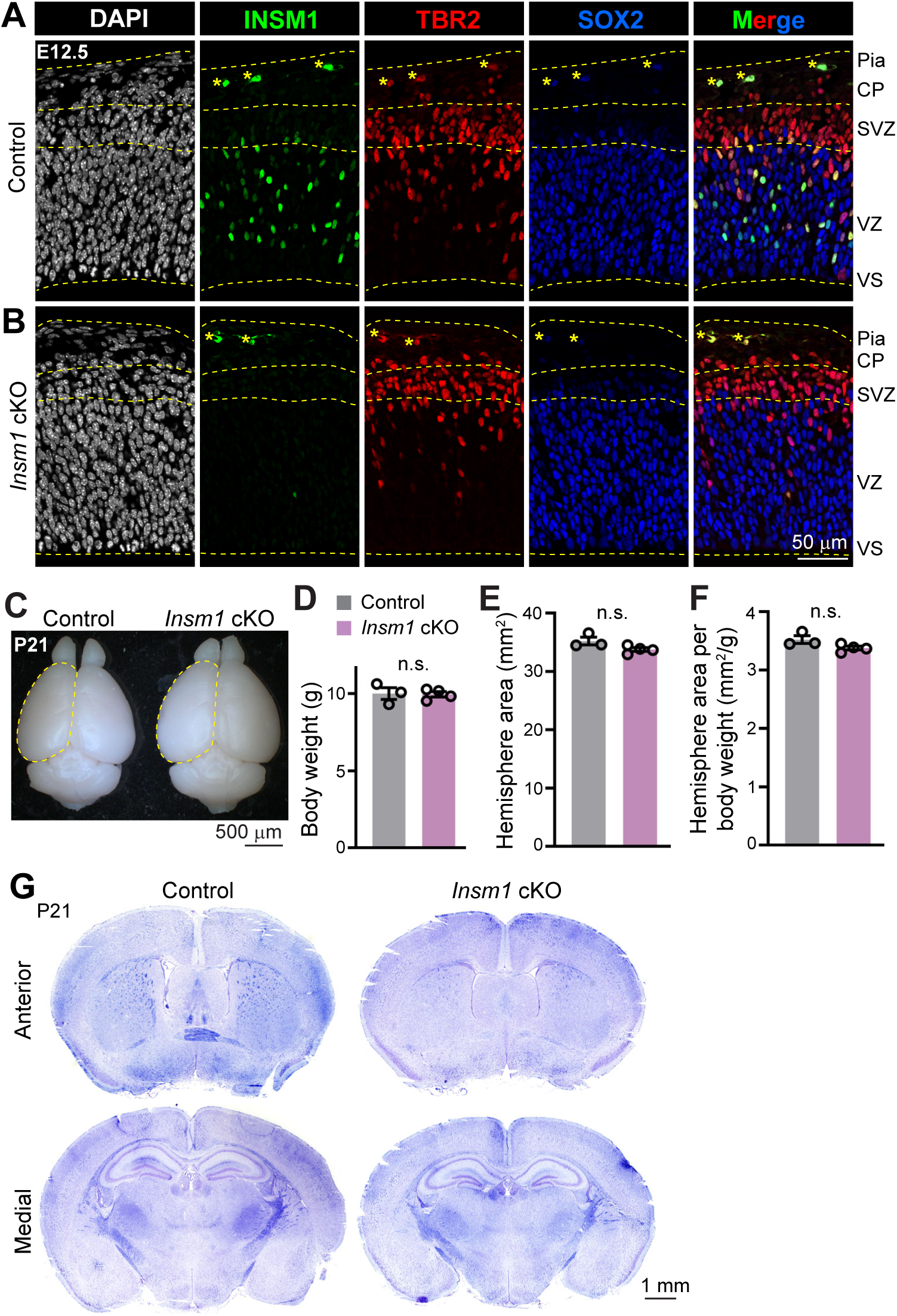
Cortical development is largely preserved in the absence of INSM1. (**A**, **B**) Neocortical regions of E12.5 control (A) and *Insm1* conditional knockout (cKO, B) brain sections co-immunostained for INSM1, the basal progenitor marker TBR2, and the neural progenitor marker SOX2. VS, ventricular surface; VZ, ventricular zone; SVZ, subventricular zone; CP, cortical plate; asterisks (*), blood cells. (**C**) Representative images of P21 control and *Insm1* cKO brains. (**D**–**F**) Quantification of the body weight (D), cerebral hemisphere area (outlined in C; E), and hemisphere area normalized to body weight (F) of P21 animals. (**G**) Nissl and Luxol Fast Blue staining of P21 brain sections at anterior and medial levels. Each data point represents an individual animal. Values are mean ± SEM. Unpaired, two-tailed *t*-test; n.s., not significant (*P* > 0.05).

These *Insm1* cKO mice were viable, fertile, and exhibit no overt behavioral abnormalities. At postnatal day (P) 21, overall brain morphology, cerebral hemisphere size, and bodyweight were indistinguishable from littermate controls (*Insm1^flox/flox^*) (Fig. 1C–F). Nissl and Luxol Fast Blue staining further revealed that cortical cytoarchitecture remained grossly normal in cKO brains (Fig. 1G).

We next quantified cortical projection neuron populations using established layer-specific markers. The numbers of FOXP2^+^ and CTIP2^+^ deep-layer neurons were significantly reduced in cKO compared to control cortices (Fig. 2A, B, and G). However, SATB2^+^ neurons, both those located in deep-layer and those in upper-layer positions, were present in comparable numbers in control and cKO brains, as were CUX1^+^ upper-layer neurons (Fig. 2C, D, and G). Thus, loss of INSM1 selectively affects the production of deep-layer, but not upper-layer, neurons.

**Figure 2.**
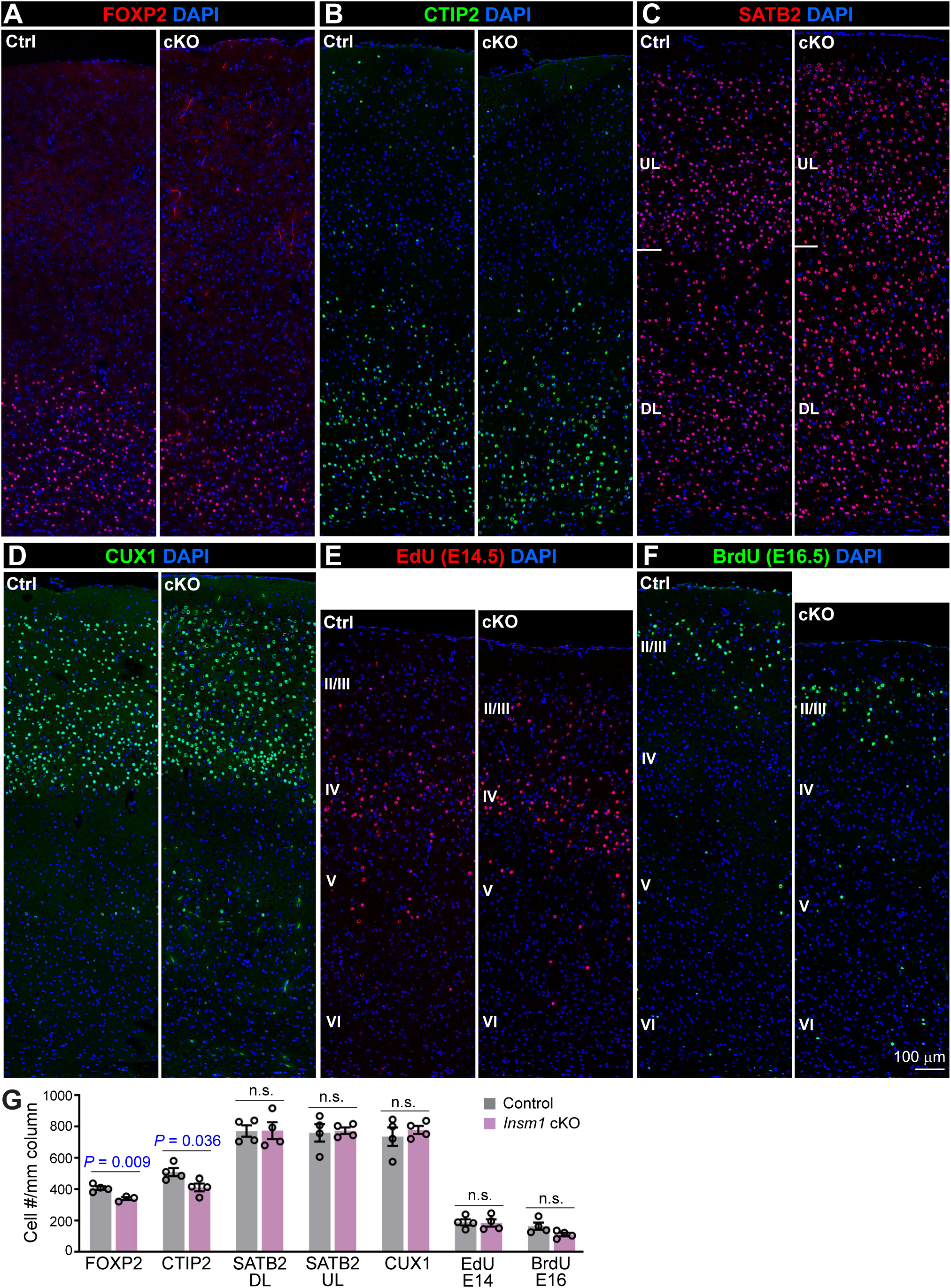
INSM1 loss reduces deep-layer neurons but spares upper-layer neurons. (**A**–**D**) Neocortical regions of P21 control (Ctrl) and *Insm1* cKO brain sections immunostained for layer-specific projection neuron markers. (**E**, **F**) Neocortical regions of P21 brains labeled by EdU (E) or BrdU (F), which were administrated at E14.5 and E16.5, respectively. (**G**) Quantification of indicated neuronal populations per mm of cortical radial column at P21. DL, deep-layer; UL, upper-layer. Each data point represents an individual animal. Values are mean ± SEM. Unpaired, two-tailed *t*-test; n.s., not significant (*P* > 0.05).

Deep- and upper-layer neurons in the mouse neocortex are born between E11.5–E14.5 and E13.5–E18.5, respectively (Caviness, 1982; Takahashi et al., 1999; Greig et al., 2013; Huilgol et al., 2025). To examine the timing of neurogenesis, we performed birth-dating experiments by administering thymidine analogs at E14.5 and E16.5. The numbers of neurons generated at these time points did not differ between control and cKO cortices (Fig. 2E–G). Moreover, the laminar positions of labeled neurons were appropriate: neurons born at E14.5 localized predominantly to layer IV, and those generated at E16.5 localized to layers II/III (Fig. 2E, F). Thus, INSM1 loss does not alter neurogenesis timing or laminar allocation.

These results contrast with an earlier report that *Insm1* deletion reduces neuron numbers across all cortical layers (Farkas et al., 2008). To reconcile this discrepancy, we performed a comprehensive analysis of the effects of INSM1 loss during mid and late cortical neurogenesis.

### INSM1 loss preserves AP and BP numbers at mid-neurogenesis but alters their proliferative behaviors

At E14.5, *Insm1* cKO cortices were grossly normal (Fig. 3A). The lateral extension of the neocortex measured along the VS (Fig. 3A, arrowheads) was similar between control and cKO brains (Fig. 3B). Immunostaining for the AP/RG marker SOX2 and the BP marker TBR2 revealed appropriate spatial distributions at this stage: in both genotypes, APs were tightly packed within the VZ, and BPs were scattered in the VZ, concentrated in the SVZ, and sparsely present in the intermediate zone (IZ)—which harbors newborn neurons en route to the cortical plate (Fig. 3C). For quantification, we defined SOX2^+^TBR2^−^ cells as APs and TBR2^+^ cells as BPs (Englund et al., 2005). AP and BP numbers were comparable between control and cKO cortices (Fig. 3D, E).

**Figure 3.**
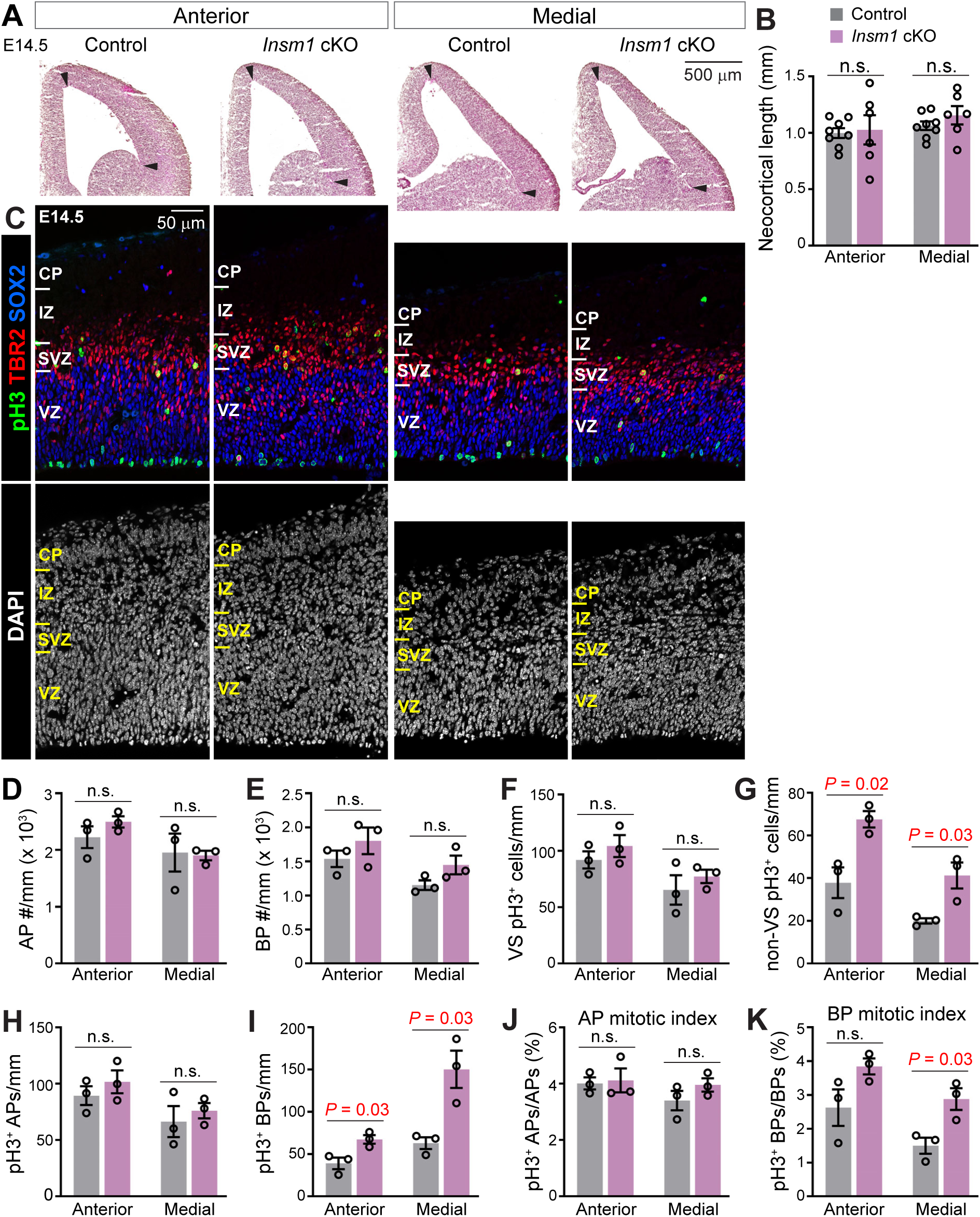
INSM1 loss preserves AP and BP numbers at mid-neurogenesis but alters BP mitosis. (**A**, **B**) Hematoxylin and eosin (H&E) staining of E14.5 brain sections at anterior and medial levels, with quantification of neocortical length. Arrowheads indicate the boundaries used for measuring neocortical length. (**C**) Neocortical regions of E14.5 brain sections co-immunostained for TBR2, SOX2, and the mitotic marker phospho-histone H3 (pH3). (**D**–**I**) Quantification of the numbers of apical progenitors (APs; SOX2^+^TBR2^−^; D), basal progenitors (BPs; TBR2^+^; E), mitotic cells located at the ventricular surface (VS; F) or away from it (non-VS; G), and mitotic APs (pH3^+^SOX2^+^TBR2^−^; H) and BPs (pH3^+^TBR2^+^; I) per mm of cortical radial column. (**J**, **K**) Mitotic index of APs (J) and BPs (K), calculated as the fraction of pH3⁺ cells within each population. Each data point represents an individual animal. Values are mean ± SEM. Unpaired, two-tailed *t*-test; n.s., not significant (*P* > 0.05).

A defining feature of APs and BPs is their mitotic location, with APs undergoing mitosis at the VS and BPs away from the VS. We quantified mitotic cells, labeled by phospho-histone H3 (pH3), separately at the VS and away from the VS (non-VS). VS-localized mitoses were similar between control and *Insm1* cKO cortices (Fig. 3F), whereas non-VS mitoses were significantly increased in cKO cortices (Fig. 3G). To validate progenitor-type identity of mitotic cells, we counted marker-defined mitoses: pH3^+^SOX2^+^TBR2^−^ cells as mitotic APs and pH3^+^TBR2^+^ cells as mitotic BPs. Consistent with the location-based analysis, the numbers of mitotic APs were unchanged (Fig. 3H), but mitotic BPs were significantly elevated in cKO cortices (Fig. 3I). The mitotic index (fraction of mitotic cells) was likewise unaffected for APs (Fig. 3J) but increased for BPs in cKO cortices (with a non-significant trend in anterior regions; Fig. 3K).

Changes in mitotic index in a population of cycling cells indicate alterations in M-phase duration relative to total cell-cycle length. For example, a higher mitotic index could result from an unchanged M-phase duration but a shorter total cell-cycle length (i.e., a faster cycling speed), or a prolonged M-phase but an unchanged total cell-cycle length. To dissect cell cycle kinetics, we used a thymidine-analog double-labeling method (Martynoga et al., 2005) to estimate S-phase length (T_S_) and total cell cycle length (T_C_) of APs and BPs, assuming all progenitor cells are actively cycling (Fig. 4A). T_S_ and T_C_ of APs and T_S_ of BPs were indistinguishable between genotypes (Fig. 4B–E). By contrast, T_C_ of BPs was significantly prolonged in *Insm1* cKO cortices (Fig. 4F). The elevated mitotic index of cKO BPs likely arises from a disproportionate lengthening of M-phase rather than accelerated cycling.

**Figure 4.**
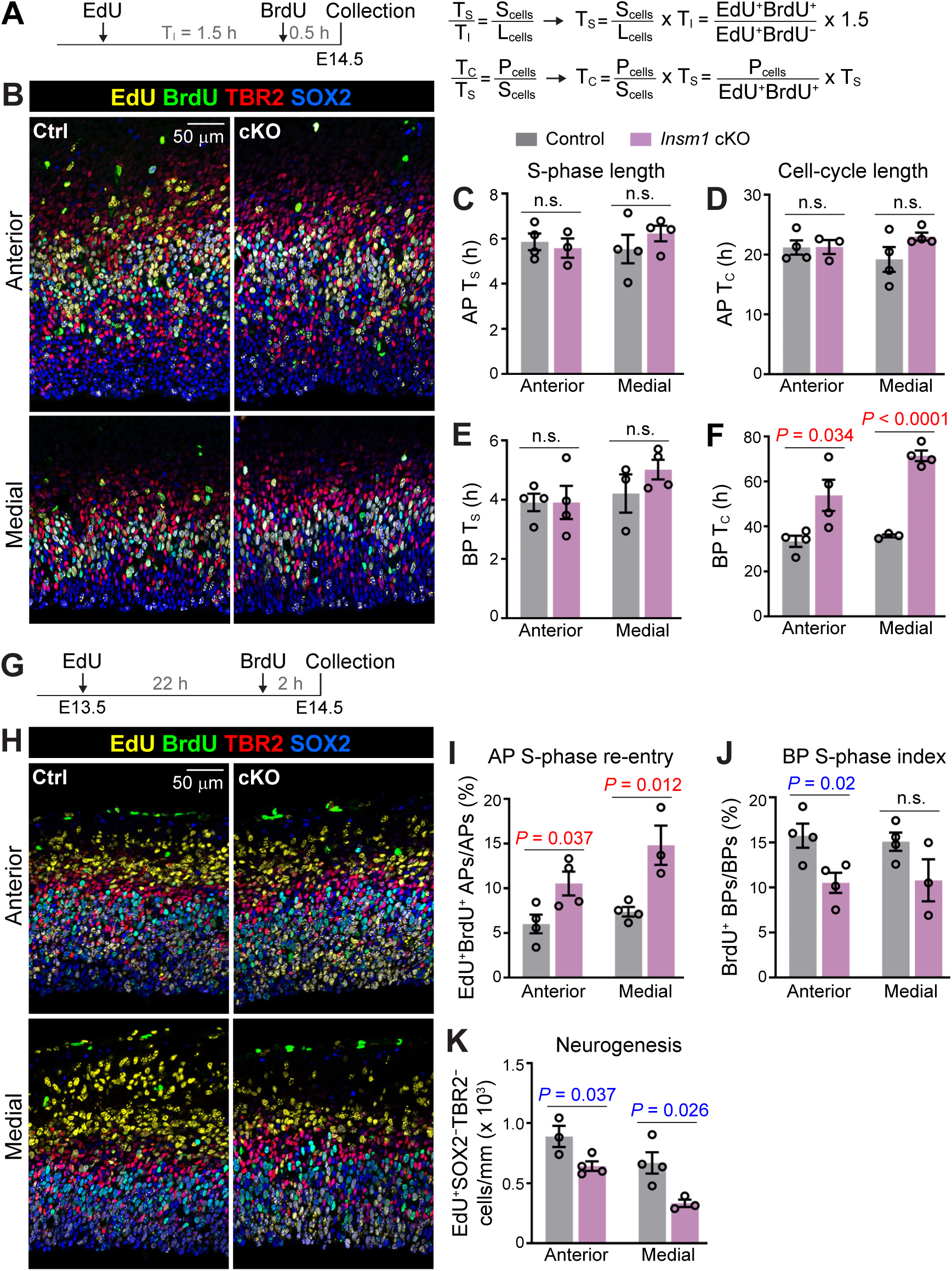
INSM1 loss alters AP and BP proliferative behaviors. (**A**) Experimental scheme and formulas of a thymidine-analog double-labeling method used to estimate S-phase length (T_S_) and total cell-cycle length (T_C_). T_I_, interval between EdU and BrdU injections (1.5 h); S_cells_, number of S-phase cells at the time of collection (EdU^+^BrdU^+^); L_cells_, leaving cells (EdU^+^BrdU^−^); P_cells_, progenitor cells. (**B**) Neocortical regions of E14.5 brain sections immunostained for cell-cycle kinetics analysis. (**C**–**F**) Quantification of cell-cycle parameters (T_S_ and T_C_) for apical progenitors (APs; SOX2^+^TBR2^−^) and basal progenitors (BPs; TBR2^+^). (**G**–**K**) Experimental scheme of EdU/BrdU double-labeling at E13.5 and E14.5 (G) and corresponding immunostaining (H) and quantification (I–K). Each data point represents an individual animal. Values are mean ± SEM. Unpaired, two-tailed *t*-test; n.s., not significant (*P* > 0.05).

To further analyze the effect of INSM1 loss on neural progenitor proliferation and differentiation, we administered EdU at E15.5 and BrdU at E16.5, and collected embryos 2 h after BrdU injection (Fig. 4G, H). The fraction of APs labeled by both EdU and BrdU, indicative of S-phase re-entry ∼22 h after the prior S-phase, was significantly higher in *Insm1* cKO cortices than in controls (Fig. 4I). Given that AP T_C_ and T_S_ were unchanged, the parsimonious explanation for the higher S-phase re-entry of cKO APs is that more APs remained as APs after the preceding division, likely by shifting from asymmetric neurogenic divisions (AP + neuron/BP) toward symmetric amplifying divisions (AP + AP). The fraction of BrdU-labeled BPs (S-phase index) was reduced in cKO (with a non-significant trend medially; Fig. 4H), consistent with the prolonged T_C_ and unchanged T_S_ of cKO BPs (Fig. 4E, F). Finally, in line with the reduced FOXP2^+^ and CTIP2^+^ deep-layer neurons at P21 (Fig. 2A, B, and G), neurons born at E13.5 (EdU^+^SOX2^−^TBR2^−^)—a stage when deep-layer neurons are being produced—were significantly decreased in cKO cortices (Fig. 4K).

Together, these data indicate that INSM1 loss slows BP cell-cycle progression and biases AP toward symmetric amplifying divisions. However, despite these changes, the loss of INSM1 does not yet have a significant impact on AP or BP numbers by mid-neurogenesis.

### INSM1 loss expands AP population and impairs BP proliferation during late neurogenesis

We next examined the consequences of INSM1 loss at E16.5, a stage when upper-layer neurogenesis is underway. At this time point, the lateral extension of the neocortex was significantly increased in *Insm1* cKO brains compared to controls (Fig. 5A, B). The VZ was noticeably thicker in cKO cortices (Fig. 5C) and AP number was significantly elevated (Fig. 5D). These findings align with our earlier conclusion that INSM1 loss enhances symmetric amplifying divisions of APs during mid-neurogenesis.

**Figure 5.**
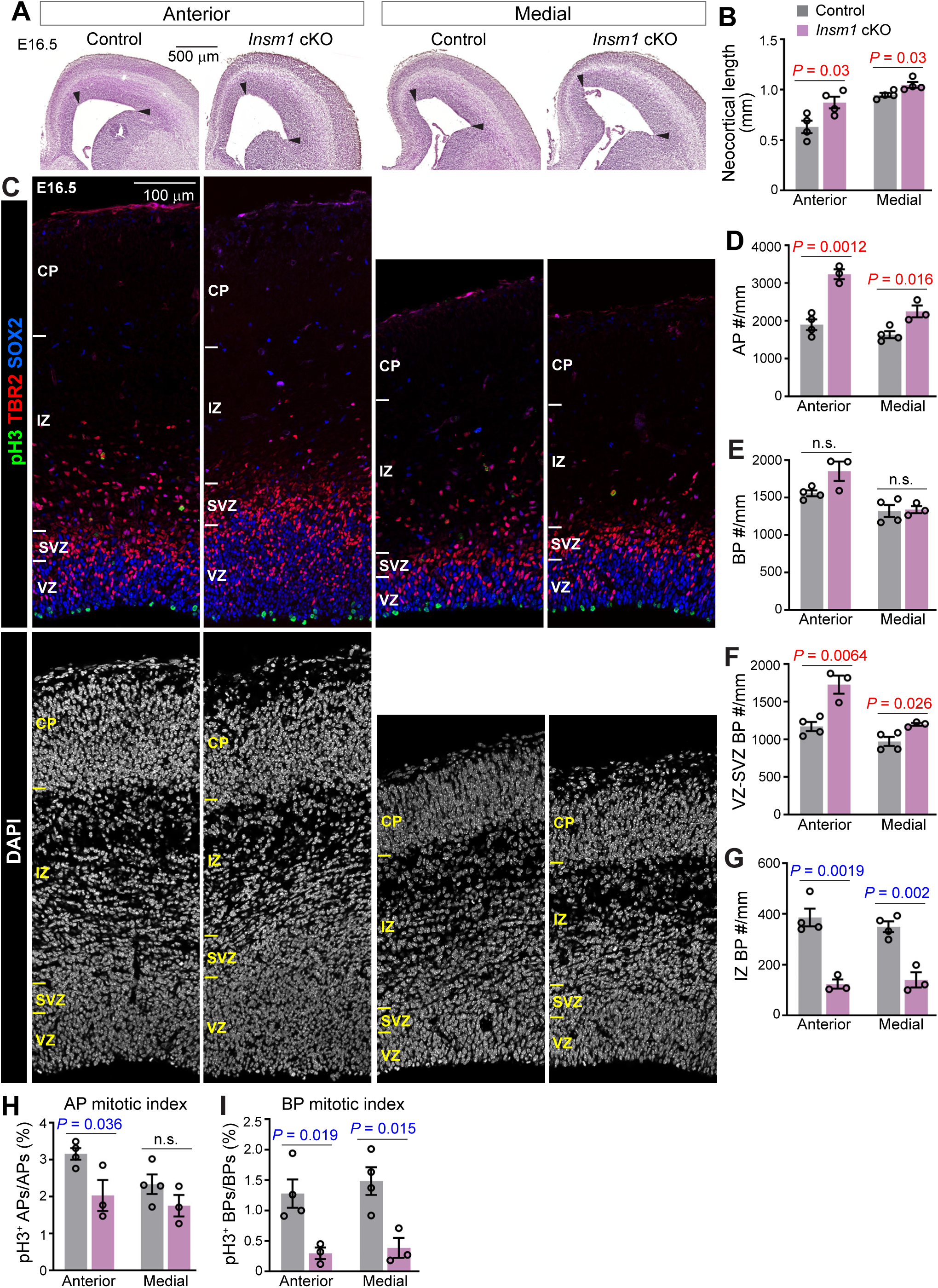
INSM1 loss expands AP population and alters BP distribution and mitosis during late neurogenesis. (**A**, **B**) H&E staining of E16.5 brain sections at anterior and medial levels, with quantification of neocortical length. Arrowheads indicate the boundaries used for measuring neocortical length. (**C**) Neocortical regions of E16.5 brain sections co-immunostained for TBR2, SOX2, and the mitotic marker phospho-histone H3 (pH3). (**D**–**I**) Quantification of the numbers of apical progenitors (APs; SOX2^+^TBR2^−^; D), basal progenitors (BPs; TBR2^+^; E), and BPs located at the ventricular and subventricular zones (VZ-SVZ; F) or the intermediate zone (IZ; G) per mm of cortical radial column. (**H**, **I**) Mitotic index of APs (H) and BPs (I), calculated as the fraction of pH3⁺ cells within each population. Each data point represents an individual animal. Values are mean ± SEM. Unpaired, two-tailed *t*-test; n.s., not significant (*P* > 0.05).

Similar to the effect at E14.5, BP number remained unchanged in cKO cortices at E16.5 (Fig. 5E). However, their spatial distribution was altered, with significantly more BPs residing in the VZ and SVZ and fewer in the IZ in cKO cortices compared with controls (Fig. 5F, G). Furthermore, the BP mitotic index was strongly reduced at E16.5 (Fig. 5I), opposite to the increased mitotic index observed at E14.5 (Fig. 3K). The AP mitotic index was decreased in anterior but not medial cortical regions (Fig. 5H).

To further assess progenitor proliferation and differentiation, we administered EdU at E15.5 and BrdU at E16.5, collecting embryos 2 h after BrdU injection (Fig. 6A, B). AP S-phase re-entry (EdU^+^BrdU^+^), which was elevated in cKO cortices at E14.5 (Fig. 4I), was no longer significantly different between control and cKO cortices at E16.5 (Fig. 6C). However, the BP S-phase index (BrdU^+^) remained significantly reduced in cKO cortices (Fig. 6E), whereas the AP S-phase index was unchanged (Fig. 6D). Because S-phase entry requires phosphorylation of the retinoblastoma protein (RB) (Massagué, 2004), we assessed phospho-RB (pRB) immunoreactivity (Fig. 6B). Consistent with the S-phase indices, the proportion of pRB^+^ APs was unchanged between control and cKO cortices (Fig. 6F), whereas that of pRB^+^ BPs was significantly reduced in cKO cortices (Fig. 6G). Notably, although the proportion of pRB^+^ BPs in the VZ was unaffected, those in the SVZ and IZ were significantly reduced in cKO cortices (Fig. 6H). Because BPs are born at the VS after AP mitotic division and progress through G1–S–G2–M phases as they migrate outward through the VZ, SVZ, and IZ, these regional deficits indicate a progressive failure of BP cell-cycle progression as they migrate away from the VS.

**Figure 6.**
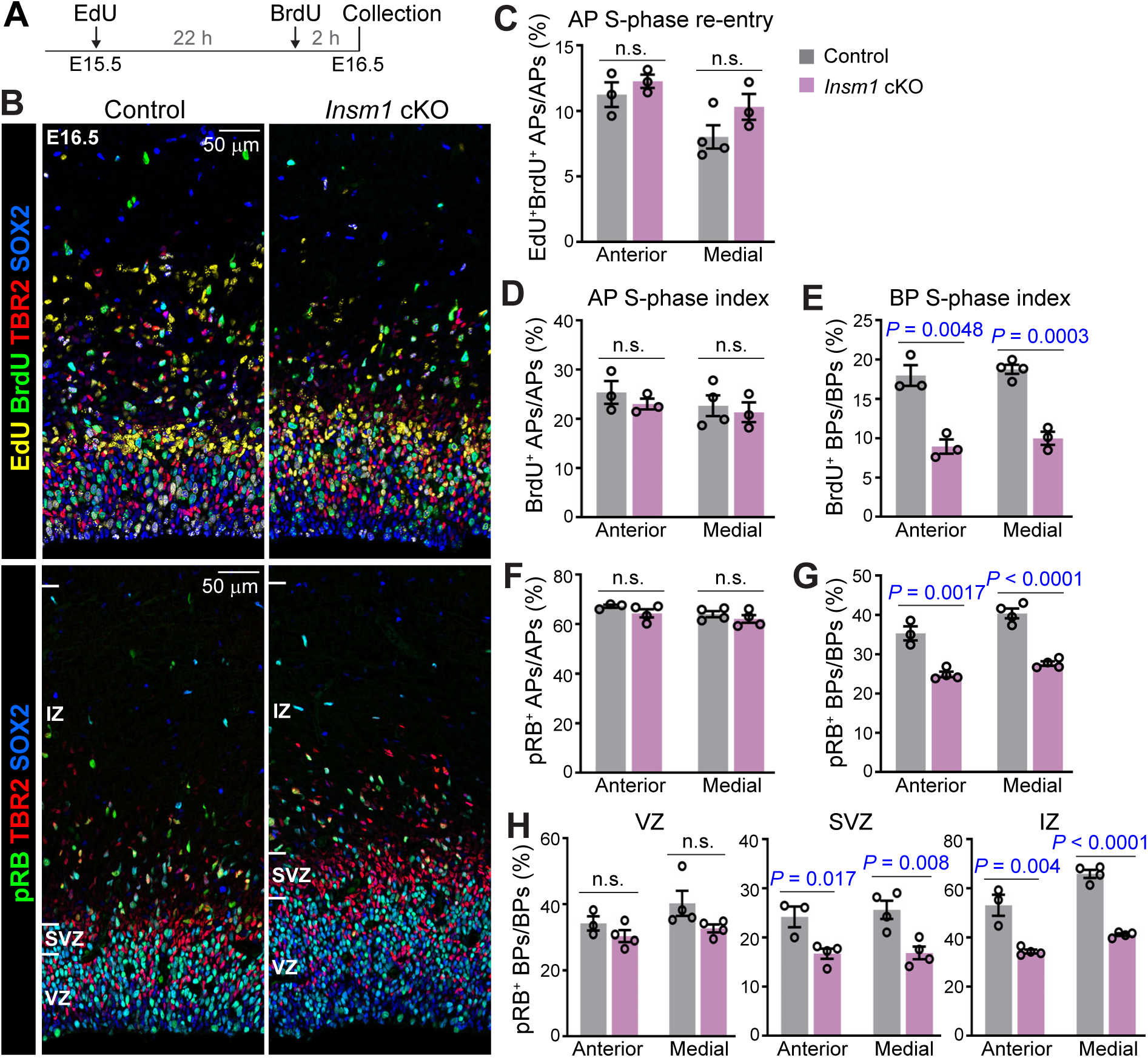
INSM1 loss impairs BP cell-cycle progression. (**A**) Experimental scheme of EdU/BrdU double-labeling at E15.5 and E16.5. (**B**) Neocortical regions of E16.5 brain sections immunostained for BrdU, TBR2, SOX2, and phospho-RB (pRB), and detected for EdU, showing the areas containing the ventricular, subventricular, and intermediate zones (VZ, SVZ, and IZ). (**C**–**E**) Quantification of S-phase re-entry in apical progenitors (APs; SOX2^+^TBR2^−^; C), and S-phase index of APs (D) and basal progenitors (BPs; TBR2^+^; E). (**F**–**H**) Fraction of pRB⁺ cells within APs (F) and BPs (G), the fraction of pRB⁺ BPs stratified by spatial location within the VZ, SVZ, and IZ (H). Each data point represents an individual animal. Values are mean ± SEM. Unpaired, two-tailed *t*-test; n.s., not significant (*P* > 0.05).

### Single-cell transcriptome analysis reveals impaired BP cell-cycle progression upon INSM1 loss

Our results thus far indicate that INSM1 is largely dispensable for BP biogenesis but is required for proper BP cell-cycle progression. To further define the impact of INSM1 loss on cortical development, we performed single-cell RNA sequencing (scRNA-seq) on E14.5 and E16.5 cortices from control and *Insm1* cKO embryos (N = 3 per genotype). We first analyzed the E14.5 control dataset. Clustering based on highly variable genes followed by UMAP visualization identified nine major cell clusters (Fig. 7A). Marker gene expression revealed that these clusters were composed of APs, BPs, immature projection neurons (iPNs), immature interneurons (*Gad2*), endothelial cells (*Igfbp7*), Cajal-Retzius cells (*Reelin*), and microglia (*Cx3cr1*) (Fig. 7C). Consistent with expected proliferative states, AP and BP clusters contained cells distributed across S, G2/M, and G0/G1 phases, whereas iPN clusters contained only G0/G1 cells (Fig. 7B). These results closely matched prior scRNA-seq analyses of wild-type E14.5 cortex (Noack et al., 2022), validating our computational workflow.

**Figure 7.**
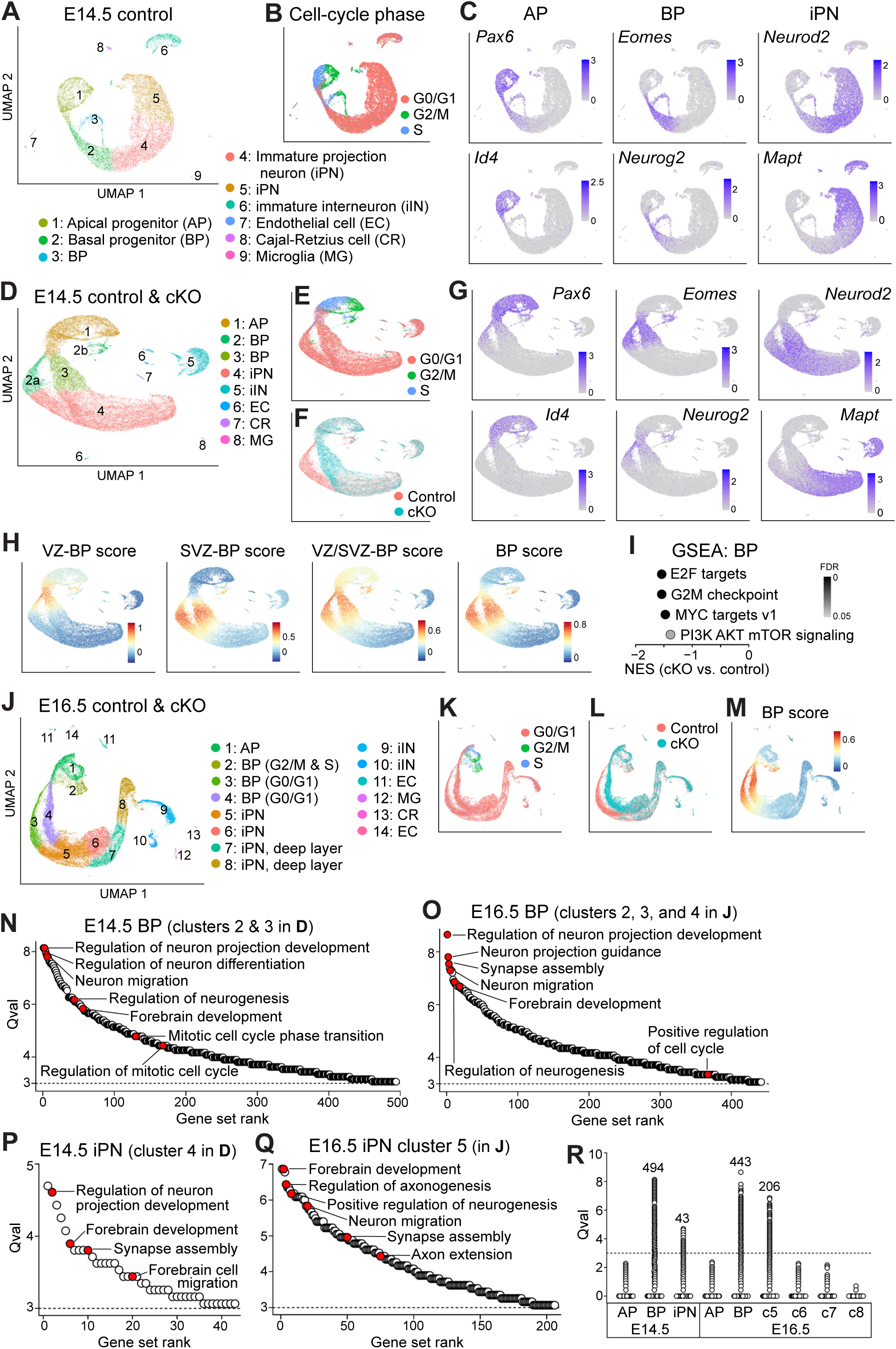
Single-cell RNA-sequencing analysis reveals BP dysregulation upon INSM1 loss. (**A**) Clustering and annotation of E14.5 control cells visualized by UMAP. *N* = 18,961 cells after quality control. (**B**) Cell-cycle phase distribution of E14.5 control cells. (**C**) Feature plots showing expression of selected marker genes used to define major cell states. (**D**) Joint clustering and annotation of E14.5 control and *Insm1* cKO cells visualized by UMAP. *N* = 40,748 cells after quality control. (**E**–**G**) Cell-cycle phase (E), genotype (F), and marker gene expression (G) for E14.5 control (*N* = 18,961) and cKO (*N* = 21,787) cells. (**H**) Signature scores for curated gene sets enriched in BPs localized predominantly to the ventricular zone (VZ-BP), subventricular zone (SVZ-BP), both zones (VZ/SVZ-BP), or all BP-enriched genes (BP). (**I**) Hallmark gene sets significantly downregulated in E14.5 *Insm1* cKO BPs compared to control BPs (cells in clusters 2 and 3 in D), identified using pseudo-bulk gene set enrichment analysis (GSEA). (**J**) Joint clustering and annotation of E16.5 control and *Insm1* cKO cells visualized by UMAP. *N* = 38,119 cells after quality control. (**K**–**M**) Cell-cycle phase (K), genotype (L), and BP score (M) for E16.5 control (*N* = 17,501) and cKO (*N* = 20,618) cells. (**N**–**Q**) Ranking of the gene sets with a Qval ≥ 3 based on single-cell pathway analysis (SCPA) comparing control and cKO cells, with selected gene sets highlighted. (**R**) Qval from SCPA comparisons of control versus cKO cells stratified by cell state or cluster. Numbers indicate the count of gene sets with a Qval ≥ 3 among the 4,132 gene sets in the GOBP collection.

We next jointly analyzed E14.5 control and cKO cells using the same methodology. APs and iPNs from both genotypes intermingled extensively and formed unified clusters, indicating minimal transcriptomic divergence in these cell states (Fig. 7D, G). In contrast, G0/G1-phase BPs segregated into two distinct clusters based on genotype, with cluster 2a containing only control BPs and cluster 3 only cKO BPs; cluster 2b contained BPs from both genotypes but predominantly S- and G2/M-phase cells (Fig. 7D–F). This selective genotype-based segregation suggests that the primary transcriptomic differences between control and cKO cortices at E14.5 arise within the BP population. To confirm that the cKO-segregated cluster (cluster 3) represented bona fide BPs, we scored cells according to curated gene sets that show (i) predominant expression in BPs located within the VZ (VZ-BP, 12 genes), (ii) predominant expression in BPs located within the SVZ (SVZ-BP, 69 genes), (iii) similar expression in BPs located in both VZ and SVZ (VZ/SVZ-BP, 55 genes), and (iv) all these BP-expressed genes (136 genes) (Supplemental Table S1); the expression patterns of these genes were validated by in situ hybridization (Bedogni and Hevner, 2021). The cKO cells in cluster 3 scored similarly to control BPs in cluster 2a (Fig. 7H), confirming that they were genuine BPs rather than cells of an aberrant cell state.

To identify the difference between cKO and control BPs (cells in clusters 2 and 3), we performed gene set enrichment analysis (GSEA) using a pseudo-bulk approach. In the Hallmark collection, only four gene sets were significantly different (FDR ≤ 0.05) between cKO and control BPs; they were all downregulated in cKO BPs and were all related to cell-cycle regulation (Fig. 7I). In the Gene Ontology Biological Process (GOBP) collection, 45 gene sets were significantly altered, 42 of which were downregulated and most were associated with cell-cycle regulation (Supplemental Table S2). These analyses strongly support the conclusion that BP cell-cycle progression is impaired in the absence of INSM1. While similar analyses of APs and iPNs also detected downregulation of some cell-cycle–related gene sets (e.g., Myc targets V1 and G2M checkpoints) in the Hallmark collection, these differences were not observed in the GOBP analysis (Supplemental Table S2); the basis for this discrepancy is unclear.

Analysis of the E16.5 scRNA-seq dataset produced similar results: while APs and iPNs from control and cKO cortices clustered together, G0/G1-phase BPs again separated into two genotype-specific clusters (clusters 3 and 4, Fig. 7J–M, Supplemental Fig. S1A, B). As at E14.5, GSEA demonstrated significant downregulation of cell-cycle related gene sets in cKO BPs and APs compared with control cells (Supplemental Table S3). Collectively, these single-cell transcriptomic analyses reveal that the most pronounced effect of INSM1 loss on cortical development occurs within the BP population, causing impaired BP cell-cycle progression at both mid- and late-neurogenesis, in agreement with our histological analyses.

### INSM1 loss perturbs neural developmental processes in BPs but does not strongly affect neuronal transcriptomes

To further investigate how INSM1 loss impacts cortical development, we applied Single Cell Pathway Analysis (SCPA), a computational tool that uses a non-parametric graph-based statistical framework to assess the multivariate, joint distribution of a set of genes in single-cell RNA-seq datasets and infer whether this gene set (pathway) is differentially regulated across conditions (Bibby et al., 2022). SCPA provides a statistic, Qval, which reflects the size of distributional change for a given pathway, and has been shown to be more sensitive than cluster-based comparisons such as pseudo-bulk GSEA. However, SCPA does not provide the direction of difference between conditions; that is, whether a pathway is up- or down-regulated. Using this approach to compare control and cKO BPs, we found that nearly 500 of the 4,132 gene sets in the GOBP collection were differentially regulated at E14.5 and at E16.5 (Qval ≥ 3). These included numerous gene sets associated with neural development, along with many associated with cell-cycle regulation (Fig. 7N, O; Supplemental Tables S4 and S5). In contrast, far fewer gene sets were dysregulated in E14.5 iPNs and E16.5 cluster 5 iPNs (which likely consisted of newborn iPNs), and they again included those associated with neural development (Fig. 7P, Q; Supplemental Tables S4 and S5). No gene set reached the Qval ≥ 3 threshold in APs or in the more mature iPN clusters at E16.5 (clusters 6, 7, and 8; Fig. 7R; Supplemental Tables S4 and S5). Taken together, these analyses indicate that INSM1 loss primarily perturbs neural developmental processes in BPs. Although some dysregulation persists into the earliest stages of neuronal differentiation, the transcriptomes of more mature neurons remain largely unaffected, underscoring the robustness of cortical development.

## Discussion

Previous studies have concluded that INSM1 “promotes the generation and expansion of basal progenitors in the developing mouse neocortex” (Farkas et al., 2008; Tavano et al., 2018). By conditionally deleting *Insm1* during mouse cortical development and analyzing its effects across APs, BPs, and neurons, we show that INSM1 is not required for the generation of cortical BPs. Instead, INSM1 is essential for proper cell-cycle progression of BPs. In the absence of INSM1, cortical BPs are defective in upregulating RB phosphorylation and entering S-phase, and exhibit downregulation of cell-cycle–related gene sets in their transcriptomes. Because the vast majority of BPs in the developing mouse neocortex undergo a single round of terminal symmetric division to generate two neurons (Haubensak et al., 2004; Miyata et al., 2004; Noctor et al., 2004; Noctor et al., 2008; Mihalas and Hevner, 2018; Huilgol et al., 2025), a defect in entering or progressing through the cell cycle likely causes many *Insm1* cKO BPs to directly differentiate into neurons without dividing, thereby reducing neuronal output from BPs. Notably, the developing cortices of reptiles and birds also contain TBR2-expressing cells, but these cells are non-proliferative and express neuronal markers (Nomura et al., 2013; Nomura et al., 2016). This raises the possibility that TBR2⁺ postmitotic neurons represent the ancestral state, and that the ability of TBR2⁺ cells to proliferate and function as BPs emerged during mammalian cortical evolution (Nomura et al., 2016). In this respect, loss of INSM1 causes mouse cortical BPs to revert toward this ancestral-like state, resembling the non-proliferative TBR2⁺ cells in sauropsid cortices.

INSM1 loss also alters the proliferative behavior of APs. At mid-neurogenesis (E14.5), more APs undergo symmetric amplifying divisions—likely at the expense of asymmetric AP–neuron divisions—resulting in a significant expansion of the AP population by late neurogenesis (E16.5). Importantly, the BP cell-cycle defect is present at both E14.5 and E16.5, whereas the change in AP division mode is detected only at E14.5. This temporal dissociation suggests that the BP defect is a direct consequence of INSM1 loss, whereas the altered AP division mode may represent a compensatory response to impaired BP proliferation. As a result, although the proliferative defect of BPs and the change in AP division mode lead to reduced production of early-born, deep-layer neurons in *Insm1* cKO cortices, the expansion of APs during late neurogenesis compensate for the reduced neurogenesis by BPs, ultimately enabling normal production of upper-layer neurons. These findings underscore the remarkable adaptability of cortical development to genetic perturbation and suggest that BP dysfunction feeds back onto APs to modulate their proliferative behavior.

BPs are widely thought to contribute to the evolutionary expansion of the mammalian cerebral cortex by increasing neuron numbers (Martínez-Cerdeño et al., 2006; Molnár et al., 2006; Cheung et al., 2007; Cheung et al., 2010; Borrell and Calegari, 2014). However, our finding that the production of upper-layer neurons remains unaffected after INSM1 loss, despite reduced neurogenesis by BPs, suggests that increasing neuron numbers may not be the sole—or even primary—function of BPs in the mouse cortex, as BPs’ contribution can be buffered by compensatory modulation of AP proliferation. Previous studies have shown that clonally related projection neurons arising from the same AP form spatially isolated radial clusters (Luskin et al., 1988; Rakic, 1988; Noctor et al., 2001; Gao et al., 2014) and preferentially develop synapses with each other over nearby non-clonally related neurons (Yu et al., 2009; Li et al., 2012; Ohtsuki et al., 2012; Yu et al., 2012). Loss of TBR2 during cortical development not only reduces BP number and the neuronal output of individual APs, but also causes lateral dispersion of clonally related projection neurons and disrupts their preferential synaptic connectivity (Lv et al., 2019), indicating that neurogenesis by BPs plays a role in microcircuit assembly. Given that upper-layer neurons in the *Insm1* cKO cortex arise from an expanded number of APs—and therefore a greater number of clones—microcircuit organization may be disrupted even though neuron numbers are preserved.

*Insm1* is broadly expressed during mammalian neural development in a pattern that generally correlates with neurogenesis; for example, within the neural tube, *Insm1* is expressed in non-surface progenitors and nascent neurons but not in apically dividing progenitors or mature neurons (Duggan et al., 2008; Farkas et al., 2008). Here we show that INSM1 promotes the neurogenic proliferation of cortical BPs, a function mirrored in delaminated otic progenitors, which are analogous to cortical BPs (Lorenzen et al., 2015). However, in the developing olfactory epithelium, INSM1 promotes the transition from early-state progenitors, including apical and transit amplifying progenitors, to terminally dividing neurogenic BPs (Rosenbaum et al., 2011). During ventral telencephalon development, INSM1 may similarly promote the progression from early- to late-state progenitors, as its loss expands the early-state, OLIG2^+^ASCL1^+^ progenitor population (Perry et al., 2025). Thus, although INSM1 promotes neurogenesis across diverse neural tissues, the precise cellular process it regulates are context specific.

INSM1 also influences neural development beyond neurogenesis. In the mammalian cochlea, *Insm1* is transiently expressed in nascent outer hair cells, and its loss causes these cells to trans-differentiate into inner hair cells (Wiwatpanit et al., 2018), demonstrating a role in neuronal subtype specification. The *C. elegans* and *Drosophila* orthologs of INSM1 regulate neuronal differentiation and neurite architecture, as well as neurogenesis (Wu et al., 2001; Yu et al., 2003; Kuzin et al., 2005; Feng et al., 2013; Froldi et al., 2015; O’Brien et al., 2017).

The molecular mechanisms by which INSM1 regulates neural development remain poorly understood. Although INSM1 contains five zinc-fingers, a recent structural study indicates that it does not bind DNA in a sequence-specific manner; instead, INSM1 is likely recruited to specific genomic loci through interactions with DNA-binding proteins (Zhou et al., 2025). In pancreatic β-cells, INSM1 interacts with NEUROD1 and FOXA2 (Jia et al., 2015), and in the ventral telencephalon, we have shown that INSM1 physically and functionally interacts with TEAD (Perry et al., 2025). Identifying the DNA-binding proteins that recruit INSM1 to its target genes across developing neural tissues and cell types will be critical for elucidating how INSM1 exerts its diverse roles during neural development.

## Competing Interest Statement

The authors declare no competing financial interests.

## Acknowledgements

This work was supported by National Institute of Health grant R01NS119760, P30CA021765 (to St. Jude Children’s Research Hospital Comprehensive Cancer Center core facilities), and American Lebanese Syrian Associated Charities (ALSAC). We thank members of the Cao lab for suggestions and technical help; Aaron Taylor and George Campbell for help in image acquisition; Abbas Shirinifard for help in image quantification; Jackie Norrie for help in single-cell RNA-seq; and Jaime García-Añoveros for sharing mouse line.

## Supplemental Material

Supplemental Figures S1. Additional data related to Figure 7.

Supplemental Table S1. Gene lists used for scRNA-seq data analysis. Related to Figure 7.

Supplemental Table S2. GSEA of scRNA-seq E14.5 datasets. Related to Figure 7.

Supplemental Table S3. GSEA of scRNA-seq E16.5 datasets. Related to Figure 7.

Supplemental Table S2. SCPA of scRNA-seq E14.5 datasets. Related to Figure 7.

Supplemental Table S3. SCPA of scRNA-seq E16.5 datasets. Related to Figure 7.

**Figure S1.**
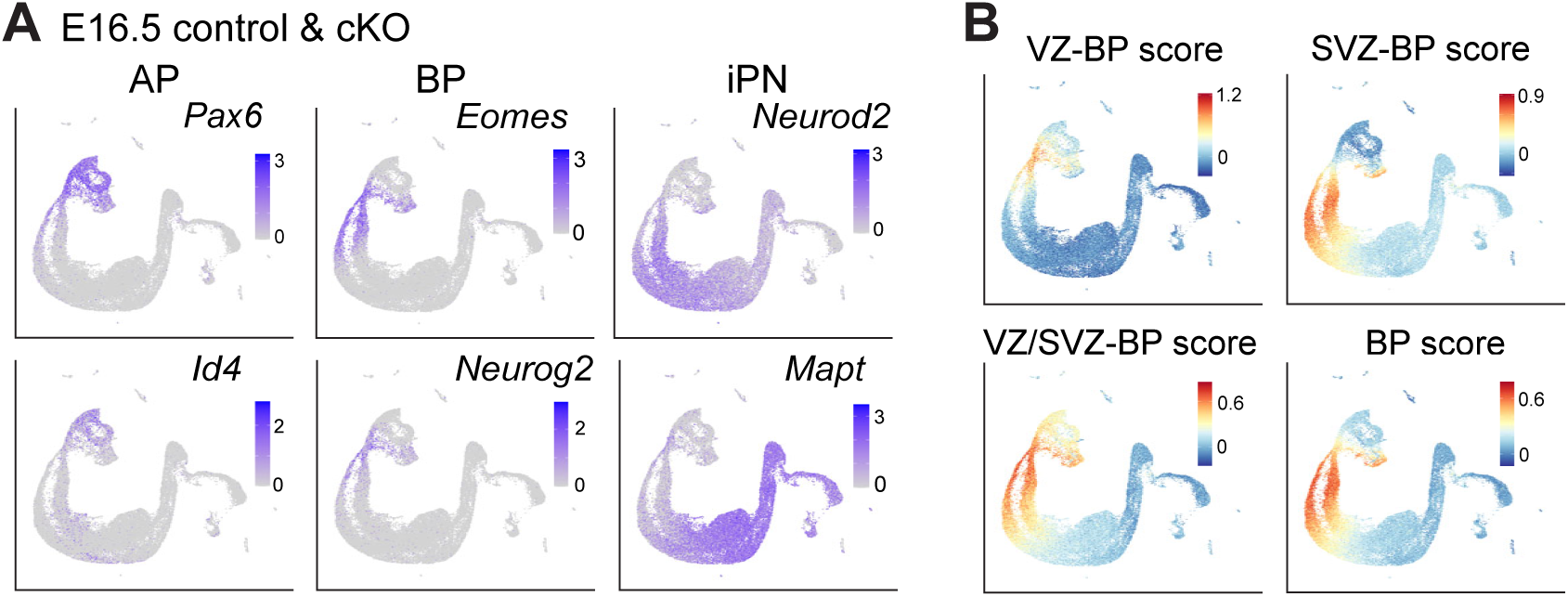
Single-cell RNA-sequencing analysis. (**A**) Feature plots of E16.5 control and *Insm1* cKO cells visualized by UMAP, showing expression of selected marker genes used to define major cell states. AP, apical progenitor; BP, basal progenitor; iPN, immature projection neuron. (**B**) Signature scores for curated gene sets enriched in BPs localized predominantly to the ventricular zone (VZ-BP), subventricular zone (SVZ-BP), both zones (VZ/SVZ-BP), or all BP-enriched genes (BP).

## Notes

### Competing Interest Statement

The authors have declared no competing interest.

## References

Angevine JB, Sidman RL (1961) Autoradiographic Study of Cell Migration during Histogenesis of Cerebral Cortex in the Mouse. Nature 192:766–768.

Bedogni F, Hevner RF (2021) Cell-Type-Specific Gene Expression in Developing Mouse Neocortex: Intermediate Progenitors Implicated in Axon Development. Front Mol Neurosci 14:686034.

Bibby JA, Agarwal D, Freiwald T, Kunz N, Merle NS, West EE, Singh P, Larochelle A, Chinian F, Mukherjee S, Afzali B, Kemper C, Zhang NR (2022) Systematic single-cell pathway analysis to characterize early T cell activation. Cell Rep 41:111697.

Borrell V, Calegari F (2014) Mechanisms of brain evolution: Regulation of neural progenitor cell diversity and cell cycle length. Neuroscience Research 86:14–24.

Breslin MB, Zhu M, Notkins AL, Lan MS (2002) Neuroendocrine differentiation factor, IA-1, is a transcriptional repressor and contains a specific DNA-binding domain: identification of consensus IA-1 binding sequence. Nucleic Acids Res 30:1038–1045.

Caviness VS (1982) Neocortical histogenesis in normal and reeler mice: A developmental study based upon [3H]thymidine autoradiography. Developmental Brain Research 4:293–302.

Cheung AFP, Pollen AA, Tavare A, Deproto J, Molnár Z (2007) Comparative aspects of cortical neurogenesis in vertebrates. In: Journal of Anatomy, 2 Edition, pp 164–176.

Cheung AFP, Kondo S, Abdel-Mannan O, Chodroff RA, Sirey TM, Bluy LE, Webber N, Deproto J, Karlen SJ, Krubitzer L, Stolp HB, Saunders NR, Molnár Z (2010) The subventricular zone is the developmental milestone of a 6-layered neocortex: Comparisons in metatherian and eutherian mammals. Cerebral Cortex 20:1071–1081.

Chiang C, Ayyanathan K (2013) Snail/Gfi-1 (SNAG) family zinc finger proteins in transcription regulation, chromatin dynamics, cell signaling, development, and disease. Cytokine Growth Factor Rev 24:123–131.

Dehay C, Huttner WB (2024) Development and evolution of the primate neocortex from a progenitor cell perspective. Development 151.

Duggan A, Madathany T, de Castro SCP, Gerrelli D, Guddati K, García-Añoveros J (2008) Transient expression of the conserved zinc finger gene INSM1 in progenitors and nascent neurons throughout embryonic and adult neurogenesis. Journal of Comparative Neurology 507:1497–1520.

Englund C, Fink A, Lau C, Pham D, Daza RA, Bulfone A, Kowalczyk T, Hevner RF (2005) Pax6, Tbr2, and Tbr1 are expressed sequentially by radial glia, intermediate progenitor cells, and postmitotic neurons in developing neocortex. The Journal of neuroscience : the official journal of the Society for Neuroscience 25:247–251.

Farkas LM, Haffner C, Giger T, Khaitovich P, Nowick K, Birchmeier C, Paabo S, Huttner WB (2008) Insulinoma-associated 1 has a panneurogenic role and promotes the generation and expansion of basal progenitors in the developing mouse neocortex. Neuron 60:40–55.

Feng G, Yi P, Yang Y, Chai Y, Tian D, Zhu Z, Liu J, Zhou F, Cheng Z, Wang X, Li W, Ou G (2013) Developmental stage-dependent transcriptional regulatory pathways control neuroblast lineage progression. Development 140:3838.

Froldi F, Szuperak M, Weng C-F, Shi W, Papenfuss AT, Cheng LY (2015) The transcription factor Nerfin-1 prevents reversion of neurons into neural stem cells. Genes & Development 29:129–143.

Gao P, Postiglione MP, Krieger TG, Hernandez L, Wang C, Han Z, Streicher C, Papusheva E, Insolera R, Chugh K, Kodish O, Huang K, Simons BD, Luo L, Hippenmeyer S, Shi SH (2014) Deterministic progenitor behavior and unitary production of neurons in the neocortex. Cell 159:775–788.

Gorski JA, Talley T, Qiu M, Puelles L, Rubenstein JL, Jones KR (2002) Cortical excitatory neurons and glia, but not GABAergic neurons, are produced in the Emx1-expressing lineage. The Journal of neuroscience : the official journal of the Society for Neuroscience 22:6309–6314.

Greig LC, Woodworth MB, Galazo MJ, Padmanabhan H, Macklis JD (2013) Molecular logic of neocortical projection neuron specification, development and diversity. Nature Reviews Neuroscience 14:755–769.

Hansen DV, Lui JH, Parker PR, Kriegstein AR (2010) Neurogenic radial glia in the outer subventricular zone of human neocortex. Nature 464:554–561.

Haubensak W, Attardo A, Denk W, Huttner WB (2004) Neurons arise in the basal neuroepithelium of the early mammalian telencephalon: a major site of neurogenesis. Proc Natl Acad Sci U S A 101:3196–3201.

Huilgol D, Levine JM, Galbavy W, Wang BS, Huang ZJ (2025) Orderly specification and precise laminar deployment of mouse cortical projection neuron types through intermediate progenitors. Dev Cell 60:1947–1957.e1943.

Huilgol D, Levine JM, Galbavy W, Wang B-S, He M, Suryanarayana SM, Huang ZJ (2023) Direct and indirect neurogenesis generate a mosaic of distinct glutamatergic projection neuron types in cerebral cortex. Neuron 111:2557–2569.e2554.

Jia S, Ivanov A, Blasevic D, Müller T, Purfürst B, Sun W, Chen W, Poy MN, Rajewsky N, Birchmeier C (2015) Insm1 cooperates with Neurod1 and Foxa2 to maintain mature pancreatic β-cell function. The EMBO journal 34:1417–1433.

Kearns N, Norville K, Frelinger JG (2023) Instant Pot for antigen retrieval: a simple, safe and economical method for use in immunohistochemistry. Biotechniques 74:237–241.

Korsunsky I, Nathan A, Millard N, Raychaudhuri S (2019) Presto scales Wilcoxon and auROC analyses to millions of observations. bioRxiv:653253.

Kuzin A, Brody T, Moore AW, Odenwald WF (2005) Nerfin-1 is required for early axon guidance decisions in the developing Drosophila CNS. Dev Biol 277:347–365.

Lan MS, Breslin MB (2009) Structure, expression, and biological function of INSM1 transcription factor in neuroendocrine differentiation. The FASEB Journal 23:2024–2033.

Li Y, Lu H, Cheng PL, Ge S, Xu H, Shi SH, Dan Y (2012) Clonally related visual cortical neurons show similar stimulus feature selectivity. Nature 486:118–121.

Lorenzen SM, Duggan A, Osipovich AB, Magnuson MA, García-Añoveros J (2015) Insm1 promotes neurogenic proliferation in delaminated otic progenitors. Mechanisms of development 138 Pt 3:233–245.

Lui JH, Hansen DV, Kriegstein AR (2011) Development and evolution of the human neocortex. Cell 146:18–36.

Luskin MB, Pearlman AL, Sanes JR (1988) Cell lineage in the cerebral cortex of the mouse studied in vivo and in vitro with a recombinant retrovirus. Neuron 1:635–647.

Lv X, Ren SQ, Zhang XJ, Shen Z, Ghosh T, Xianyu A, Gao P, Li Z, Lin S, Yu Y, Zhang Q, Groszer M, Shi SH (2019) TBR2 coordinates neurogenesis expansion and precise microcircuit organization via Protocadherin 19 in the mammalian cortex. Nat Commun 10:3946.

Martínez-Cerdeño V, Noctor SC, Kriegstein AR (2006) The role of intermediate progenitor cells in the evolutionary expansion of the cerebral cortex. Cerebral cortex (New York, NY : 1991) 16 Suppl 1:i152–161.

Martynoga B, Morrison H, Price DJ, Mason JO (2005) Foxg1 is required for specification of ventral telencephalon and region-specific regulation of dorsal telencephalic precursor proliferation and apoptosis. Developmental biology 283:113–127.

Massagué J (2004) G1 cell-cycle control and cancer. Nature 432:298–306.

Mihalas AB, Hevner RF (2018) Clonal analysis reveals laminar fate multipotency and daughter cell apoptosis of mouse cortical intermediate progenitors. Development 145.

Miyata T, Kawaguchi A, Saito K, Kawano M, Muto T, Ogawa M (2004) Asymmetric production of surface-dividing and non-surface-dividing cortical progenitor cells. Development 131:3133–3145.

Molnár Z, Métin C, Stoykova A, Tarabykin V, Price DJ, Francis F, Meyer G, Dehay C, Kennedy H (2006) Comparative aspects of cerebral cortical development. European Journal of Neuroscience 23:921–934.

Mootha VK et al. (2003) PGC-1α-responsive genes involved in oxidative phosphorylation are coordinately downregulated in human diabetes. Nature Genetics 34:267–273.

Noack F, Vangelisti S, Raffl G, Carido M, Diwakar J, Chong F, Bonev B (2022) Multimodal profiling of the transcriptional regulatory landscape of the developing mouse cortex identifies Neurog2 as a key epigenome remodeler. Nat Neurosci 25:154–167.

Noctor SC, Martínez-Cerdeño V, Kriegstein AR (2008) Distinct behaviors of neural stem and progenitor cells underlie cortical neurogenesis. The Journal of comparative neurology 508:28–44.

Noctor SC, Martínez-Cerdeño V, Ivic L, Kriegstein AR (2004) Cortical neurons arise in symmetric and asymmetric division zones and migrate through specific phases. Nat Neurosci 7:136–144.

Noctor SC, Flint AC, Weissman TA, Dammerman RS, Kriegstein AR (2001) Neurons derived from radial glial cells establish radial units in neocortex. Nature 409:714–720.

Nomura T, Gotoh H, Ono K (2013) Changes in the regulation of cortical neurogenesis contribute to encephalization during amniote brain evolution. Nature Communications 4:2206.

Nomura T, Ohtaka-Maruyama C, Yamashita W, Wakamatsu Y, Murakami Y, Calegari F, Suzuki K, Gotoh H, Ono K (2016) The evolution of basal progenitors in the developing non-mammalian brain. Development 143:66–74.

O’Brien BMJ, Palumbos SD, Novakovic M, Shang X, Sundararajan L, Miller DM (2017) Separate transcriptionally regulated pathways specify distinct classes of sister dendrites in a nociceptive neuron. Developmental Biology 432:248–257.

Ohtsuki G, Nishiyama M, Yoshida T, Murakami T, Histed M, Lois C, Ohki K (2012) Similarity of visual selectivity among clonally related neurons in visual cortex. Neuron 75:65–72.

Perry CH, Lavado A, Thulabandu V, Ramirez C, Paré J, Dixit R, Mishra A, Yang J, Yu J, Cao X (2025) TEAD switches interacting partners along neural progenitor lineage progression to execute distinct functions. Genes Dev 39:849–867.

Rakic P (1974) Neurons in rhesus monkey visual cortex: systematic relation between time of origin and eventual disposition. Science 183:425–427.

Rakic P (1988) Specification of cerebral cortical areas. Science 241:170–176.

Rosenbaum JN, Duggan A, García-Añoveros J (2011) Insm1 promotes the transition of olfactory progenitors from apical and proliferative to basal, terminally dividing and neuronogenic. Neural development 6:6.

Subramanian A, Tamayo P, Mootha VK, Mukherjee S, Ebert BL, Gillette MA, Paulovich A, Pomeroy SL, Golub TR, Lander ES, Mesirov JP (2005) Gene set enrichment analysis: a knowledge-based approach for interpreting genome-wide expression profiles. Proc Natl Acad Sci U S A 102:15545–15550.

Takahashi T, Goto T, Miyama S, Nowakowski RS, Caviness VS (1999) Sequence of Neuron Origin and Neocortical Laminar Fate: Relation to Cell Cycle of Origin in the Developing Murine Cerebral Wall. The Journal of Neuroscience 19:10357–10371.

Tavano S, Taverna E, Kalebic N, Haffner C, Namba T, Dahl A, Wilsch-Bräuninger M, Paridaen JTML, Huttner WB (2018) Insm1 Induces Neural Progenitor Delamination in Developing Neocortex via Downregulation of the Adherens Junction Belt-Specific Protein Plekha7. Neuron 97:1299–1314.e1298.

Taverna E, Götz M, Huttner WB (2014) The cell biology of neurogenesis: toward an understanding of the development and evolution of the neocortex. Annu Rev Cell Dev Biol 30:465–502.

Thor S (2024) Indirect neurogenesis in space and time. Nat Rev Neurosci 25:519–534.

Wiwatpanit T, Lorenzen SM, Cantú JA, Foo CZ, Hogan AK, Márquez F, Clancy JC, Schipma MJ, Cheatham MA, Duggan A, García-Añoveros J (2018) Trans-differentiation of outer hair cells into inner hair cells in the absence of INSM1. Nature 563:691–695.

Wu J, Duggan A, Chalfie M (2001) Inhibition of touch cell fate by egl-44 and egl-46 in C. elegans. Genes & Development 15:789–802.

Yu H, Prétôt RF, Bürglin TR, Sternberg PW (2003) Distinct roles of transcription factors EGL-46 and DAF-19 in specifying the functionality of a polycystin-expressing sensory neuron necessary for C. elegans male vulva location behavior. Development 130:5217–5227.

Yu YC, Bultje RS, Wang X, Shi SH (2009) Specific synapses develop preferentially among sister excitatory neurons in the neocortex. Nature 458:501–504.

Yu YC, He S, Chen S, Fu Y, Brown KN, Yao XH, Ma J, Gao KP, Sosinsky GE, Huang K, Shi SH (2012) Preferential electrical coupling regulates neocortical lineage-dependent microcircuit assembly. Nature 486:113–117.

Zhou H, He X, Xiong Y, Gong Y, Zhang Y, Li S, Hu R, Li Y, Zhang X, Zhou X, Zhu J, Yang Y, Liu M (2025) Structural insights into a highly flexible zinc finger module unravel INSM1 function in transcription regulation. Nat Commun 16:2162.

